# 53BP1 nuclear body-marked replication stress in a human mammary cell model of BRCA2 deficiency

**DOI:** 10.1101/462119

**Authors:** Weiran Feng, Maria Jasin

**Affiliations:** Developmental Biology Program, Memorial Sloan Kettering Cancer Center, 1275 York Avenue, New York, NY 10065, USA; Louis V. Gerstner Jr. Graduate School of Biomedical Sciences, Memorial Sloan Kettering Cancer Center, 1275 York Avenue, New York, NY 10065, USA

**Keywords:** BRCA2, replication stress, 53BP1 nuclear body, replication fork protection, replication fork reversal, breast cancer, allele specific editing, CRISPR-Cas9

## Abstract

BRCA2 deficiency causes genome instability and breast and ovarian cancer predisposition, but also paradoxically promotes cell lethality. The nature of the acute, detrimental consequences of BRCA2 loss is not fully understood. We recently generated *BRCA2* conditional models from a non-transformed human mammary cell line, through allele-specific gene targeting using CRISPR-Cas9, which we now describe. With these models, we discovered that BRCA2 deficiency triggers a DNA under replication-53BP1 nuclear body formation-G1 arrest axis due to homologous recombination defects. In this Extra-view, we extend these findings to show that 53BP1 nuclear bodies are spatially linked with downstream p53 activation. Replication stress that leads to common fragile site expression to induce 53BP1 nuclear body formation is aggravated by the loss of BRCA2. Additionally, replication stress that does not selectively lead to common fragile site expression is also able to induce 53BP1 nuclear body formation in BRCA2-deficient cells, indicating lesions form more globally throughout the genome. Furthermore, compromising replication fork reversal by SMARCAL1 depletion restores replication fork protection but does not diminish the high replication stress in the BRCA2-deficient cells, further emphasizing that fork protection plays a minor role in these cells. These results further elucidate the causes and consequences of replication stresses in the face of BRCA2 inactivation, providing insight into the barriers that need to be overcome for cells to become tumorigenic.

## Introduction

*BRCA2*, along with *BRCA1*, was identified as tumor suppressor genes of hereditary breast and ovarian cancers more than two decades ago [1, 2, 3]. With the widespread sequencing of tumor genomes, somatic as well as germline *BRCA2* mutations have been associated with both the onset and advanced stages of several tumor types [4, 5, 6]. Both BRCA1 and BRCA2 are critical in preventing genome instability, a hallmark of cancer. We now know that they play roles in at least two processes essential for genome integrity maintenance: homologous recombination (HR) to repair lethal DNA lesions, including double-strand breaks, and replication fork protection (FP) to prevent nascent strand degradation at stalled replication forks [7].

While sharing many key players, including BRCA2, HR and FP are both genetically and functionally separable processes [8]. Our mechanistic understanding of FP has recently advanced by the discovery of the role of replication fork reversal, where a fork regresses to form a four-way “chicken foot”-like structure [9]. In a FP-deficient background, reversed forks serve as the substrate for subsequent nascent strand degradation by nucleases [10, 11, 12, 13]. An HR defect in *BRCA1/2*-mutated cancers was initially leveraged to develop synthetic-lethality-based therapeutic strategies, while subsequent restoration of HR and/or possibly FP in tumors confers resistance [7]. The relative contribution of HR, FP, and now fork reversal to genome integrity is complex and may be contingent upon the cellular context [8].

Modeling BRCA2 deficiency in a non-transformed background may provide insight into the early events that initiate *BRCA2*-mutated breast cancer formation. To this end, we recently leveraged the CRISPR (Clustered Regularly Interspaced Short Palindromic Repeats)-Cas9 tools to generate two *BRCA2* conditional models in MCF10A cells [14], a non-transformed human mammary epithelial cell line [15]. In the process, we also developed a CRISPR-Cas9-based approach for allele-specific gene targeting at the *BRCA2* locus.

With these *BRCA2* conditional models, we demonstrated that BRCA2 deficiency leads to replication stress and DNA under replication that in turn causes abnormalities during mitosis and 53BP1 nuclear body formation in the subsequent G1 phase, accompanied by a p53-mediated G1 arrest and ultimately cell lethality [14]. In dissecting the relative contribution of the HR and FP pathways, we provided evidence that cell viability and replication stress suppression are mainly mediated by the HR function of BRCA2, but not FP, in these non-transformed human mammary epithelial cells [14]. Here, we further explore the consequences of BRCA2 deficiency, with a focus on 53BP1 nuclear bodies, in particular their possible upstream triggers and downstream link with p53 activation. The role of the fork reversal protein SMARCAL1 and its functional interaction with BRCA2 is also examined.

## Results and discussion

### Allele-specific editing of the *BRCA2* locus

While generating the two BRCA2 conditional models in MCF10A cells [14], CRISPR-Cas9-facilitated gene targeting was employed to achieve the desired editing outcome. One of the *BRCA2* conditional systems we described involved first targeting two loxP sites to flank exons 3 and 4 of one allele (*fl)*; the remaining *WT* allele was targeted using a promoterless selectable marker inserted at the *BRCA2* start codon (Hyg-targeted) (**Fig. S1**) [14]. However, the start codon resides in exon 2, which is present in both the *fl* and *WT* alleles. Therefore, one technical challenge we anticipated was to specifically target the *WT* allele with the Hyg-targeting vector, while leaving exon 2 of the *fl* allele intact (**Fig. 1A**).

**Figure 1.**
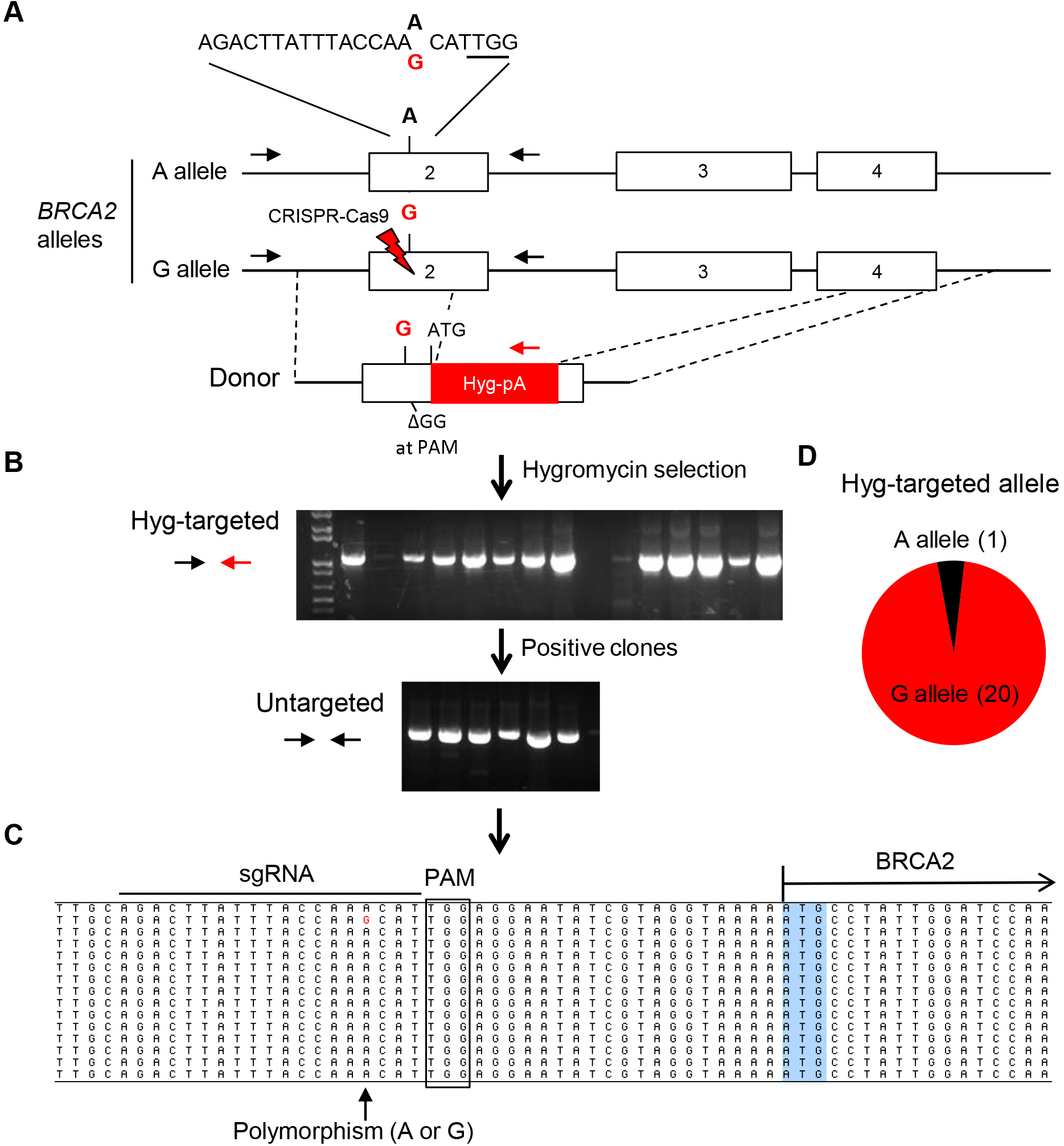
Presence of a SNP at the BRCA2 sgRNA site facilitates allele-specific gene editing. (A) Schematic for specifically targeting one *BRCA2* allele in MCF10A cells with a Hyg cassette. The BRCA2 sequence containing the A/G SNP is shown; the sgRNA sequence specifically targets the G allele (PAM sequence underlined). The donor fragment for generating the Hyg-targeted *BRCA2* null allele also contains the G SNP but it is close by to a disrupted PAM sequence (∆GG). Primer sites for PCR are indicated. Hyg, promoterless hygromycin-resistance gene; lightning bolt, CRISPR-Cas9 cleavage site. (B) Genotyping PCR using primer sets as indicated in (A). After transfection with plasmids expressing Cas9, the sgRNA, and the donor fragment, hygromycin resistant clones were subjected to a PCR to identify those that underwent successful Hyg gene targeting. PCR specific to the untargeted allele was then performed in the positive clones. A positive product is indicative of monoallelic targeting. (C) Sequences of the untargeted-allele-specific PCR products obtained in (B). Positions of the sgRNA recognition region, PAM site, SNP site, and BRCA2 coding region are indicated. (D) Hyg targeting overwhelmingly occurred at the G allele. The number of clones for each genotype at the Hyg-targeted allele is shown in parentheses, as inferred from results in (C).

To this end, we developed a method to achieve allele-specific targeting, which we benchmarked in *BRCA2^+/+^* cells to demonstrate its generality. We first sequenced the *BRCA2* exon 2 region and identified a single nucleotide polymorphism (SNP). The sgRNA was then designed specifically for the allele containing the G nucleotide SNP (“G allele”) residing towards the 3’ end of the sgRNA sequence (**Fig. 1A**), i.e., the seed region of the sgRNA that is sensitive to mismatches [16, 17, 18]. The post-transfection clones showing Hyg resistance were screened by a PCR specific for the Hyg cassette correctly integrated at the *BRCA2* locus (**Fig. 1B**). To characterize the other allele in the positive clones, PCR was performed with a reverse primer specific to an untargeted *BRCA2* allele. All clones tested exhibited a positive PCR product, indicating monoallelic gene targeting with the Hyg cassette (**Fig. 1B**). Sequencing of these PCR products revealed a strong bias (95.2%, 19/20) towards the allele with the A nucleotide SNP (“A allele”), such that Hyg targeting predominantly occurred at the G allele (**Fig. 1C, D**). Therefore, polymorphism of even a single nucleotide at the sgRNA site can strongly bias targeting towards one specific allele. We then applied this strategy to our *BRCA2^fl/+^* cells, in this case targeting the A allele with the Hyg cassette using an A allele-specific sgRNA, as the prior *fl* targeting turned out to occur to the G allele (**Fig. S1**).

Achieving allele discrimination provides advantages for a variety of applications. For example, selectively disrupting or correcting dominant disease-causing alleles has potential clinical value. Accomplishing allele discrimination can also benefit basic research, such as achieving allele-specific imaging or studying one particular allele of interest, e.g. the maternal vs. the paternal allele. While alleles can be effectively distinguished via CRISPR approaches in cases where a unique PAM is present in one allele [19, 20], discrimination between alleles with single nucleotide mismatches within the sgRNA site is conceptually challenging given the promiscuity of sgRNA recognition [16, 17, 18]. In this regard, previous studies have shown promise in utilizing SNPs in the sgRNA target regions to achieve allele-specific mutagenesis [21, 22, 23, 24, 25]. Our results now extend allele-specific applications to HR-mediated gene targeting. Notably, allele selectivity can be further improved with a truncated sgRNA design [25], since it displays an even lower tolerance to mismatches [26]. These efforts have substantially broadened the scope of allele-specific applications to achieve various desired outcomes.

### Replication stress-vulnerable sites in BRCA2-deficient cells

One previously unappreciated consequence of BRCA2 deficiency recently uncovered by others and us is the manifestation of S/G2 repair defects in later cell cycle stages, resulting in 53BP1 nuclear body formation in the subsequent G1 phase [14, 27]. What remains mysterious is the source of the replication stress. Understanding how 53BP1 nuclear bodies are formed is key to addressing this question.

53BP1 nuclear bodies were initially reported to mark common fragile sites (CFSs). Treatment with low-dose aphidicolin (APH) is a commonly used method to specifically induce CFS instability [28]. It acts by perturbing DNA replication (polymerase α) without substantially affecting cell cycle progression, leading to breakage at the fragile sites in mitosis, termed CFS expression. The same treatment was found to massively stimulate formation of 53BP1 nuclear bodies at CFSs [29, 30]. By contrast, high doses of another replication inhibitor, hydroxyurea (HU), which does not display CFS selectivity, presumably due to a different a mode of action (in part, nucleotide pool depletion) [28], fails to induce 53BP1 nuclear bodies [29]. Thus, in genetically unmodified cells, 53BP1 nuclear body formation appears to be associated with CFS expression.

To explore the sources of 53BP1 nuclear bodies arising from BRCA2 deficiency, we first benchmarked how control MCF10A cells respond to different replication stresses induced by APH and HU. BRCA2-proficient cells displayed ≥5-fold increase in these nuclear bodies with low doses of APH (**Fig. 2A**), consistent with a recent report [31]. 53BP1 nuclear bodies were somewhat increased with HU treatment [14], but the level did not reach that seen with APH and this trend was not statistically significant (**Fig. 2B**). These results are consistent with previous findings [29], showing that conditions that induce CFS expression (low-dose APH) lead to 53BP1 nuclear body formation, while HU exerts a minimal response at the dosage used.

**Figure 2.**
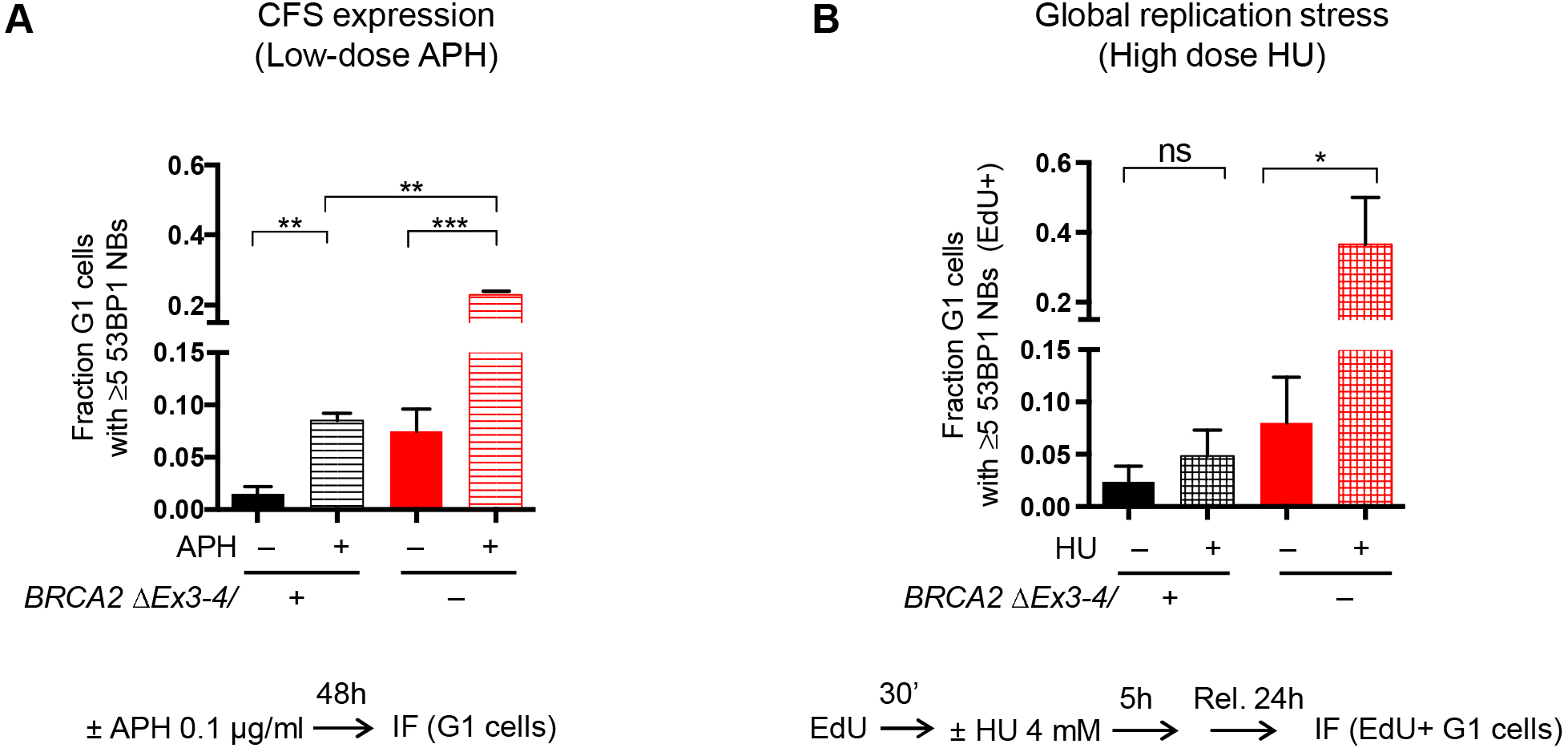
BRCA2 suppresses replication stress induced by both low-dose APH, which selectively triggers CFS expression, and HU, a more global inducer of replication stress. (A) BRCA2 deficiency further exacerbates 53BP1 nuclear body formation in cells treated with low-dose APH. Population of *BRCA2^fl/–^* MCF10A cells after acute lentiviral transduction of self-deleting Cre recombinase [57], hereafter named *BRCA2^∆Ex3-4/–^* [14], were analyzed for 53BP1 nuclear body formation induced by continuous exposure to low-dose APH, as in the schematic at bottom. The fraction of G1 cells (cyclinA-negative with 2N DNA content) containing ≥ 5 53BP1 nuclear bodies (NBs) as determined by immunofluorescence (IF) is shown. (B) BRCA2 deficiency leads to 53BP1 nuclear body formation in cells treated with HU. Cells were analyzed for HU-induced 53BP1 nuclear body formation, as in the schematic at bottom. EdU was used to label S phase cells at the time of HU treatment, which was followed by a release of the cells into the next G1 phase. EdU-positive, G1 cells (cyclinA-negative with 2N DNA content) were analyzed by IF. Note that unlike low-dose APH in (A), HU treatment dramatically induced 53BP1 nuclear body formation specifically in BRCA2-deficient, but not control cells. (Adapted from [14] under a Creative Commons license http://creativecommons.org/licenses/by/4.0/.) Error bars in this figure represent one standard deviation from the mean (s.d.). n≥2. ns, not significant; ^*^, p<0.05; ^**^, p<0.01; ^***^, p<0.001 (unpaired two-tailed t test).

Notably, upon BRCA2 disruption, the level of 53BP1 nuclear bodies was dramatically elevated not only by HU [14], but also by APH (**Fig. 2**). Interestingly, BRCA2-deficient cells displayed a level of spontaneous 53BP1 nuclear bodies that was comparable to that of BRCA2-proficient cells treated with low-dose APH (**Fig. 2A**), which provides an estimation of the level of spontaneous replication stress in these BRCA2-disrupted cells. Thus, BRCA2 suppresses replication stress that perturbs CFSs as well as other genomic sequences. This stress in turn leads to 53BP1 nuclear body formation when BRCA2 function is compromised.

We have previously shown that HU-induced replication stress is relieved by the HR function of BRCA2 [14]. Thus, HR disruption leads to fragility of genomic regions with distinct properties from those of CFSs upon HU. These properties are reminiscent of the recently discovered early replication fragile sites (ERFSs), which are characterized as regions bound by a set of DNA repair proteins, including the HR protein BRCA1, and prone to breakage in response to HU but not low-dose APH [32]. Since BRCA2-deficient cells display a high level of replication stress under both HU and low-dose APH treatment, multiple processes likely contribute to the replication stress seen in these cells. For example, BRCA2 prevents R-loop accumulation [33, 34] and maintains the integrity of telomeres [35] and G-quadruplex-forming sites [36]. Presumably these regions are more susceptible to replication stress and become fragile upon BRCA2 loss. It would be interesting for future studies to determine the full set of genomic regions that are particularly unstable upon BRCA2 deficiency and uncover the mechanism(s) of DNA fragility.

### Link between 53BP1 nuclear bodies and p53 activation

53BP1 nuclear bodies form in G1, the cell cycle stage at which BRCA2-deficient cells arrest due to p53 activation [14]. To further characterize the link between 53BP1 nuclear bodies and p53 activation in BRCA2-deficient cells, we performed image analysis to determine whether the two proteins colocalize. Indeed, a substantial fraction (~60%) of 53BP1 nuclear bodies contained p53 and its active, phosphorylated form, p53-pS15 (**Fig. 3A,B,D,E**). This colocalization occurred irrespective of BRCA2 status, although we observed a slight, but reproducible increase in the frequency of nuclear bodies with p53 signals in BRCA2-deficient cells (**Fig. 3B,E**), possibly reflecting a positive feedback regulation of p53 levels in stressed conditions [37]. However, as a consequence of increased nuclear body abundance, BRCA2 deficiency led to a marked elevation of both p53- and p53-pS15-positive 53BP1 nuclear bodies in G1 phase (**Fig. 3C,F**). Indeed, BRCA2 disruption is accompanied with an increase in spontaneous p53 levels in various systems [14, 38, 39]. Importantly, p53 disruption did not affect 53BP1 nuclear body formation (**Fig. 3G**), suggesting that p53 functions downstream of the DNA lesions marked by 53BP1. Together, these results establish that 53BP1 nuclear bodies are spatially linked to p53 activation.

**Figure 3.**
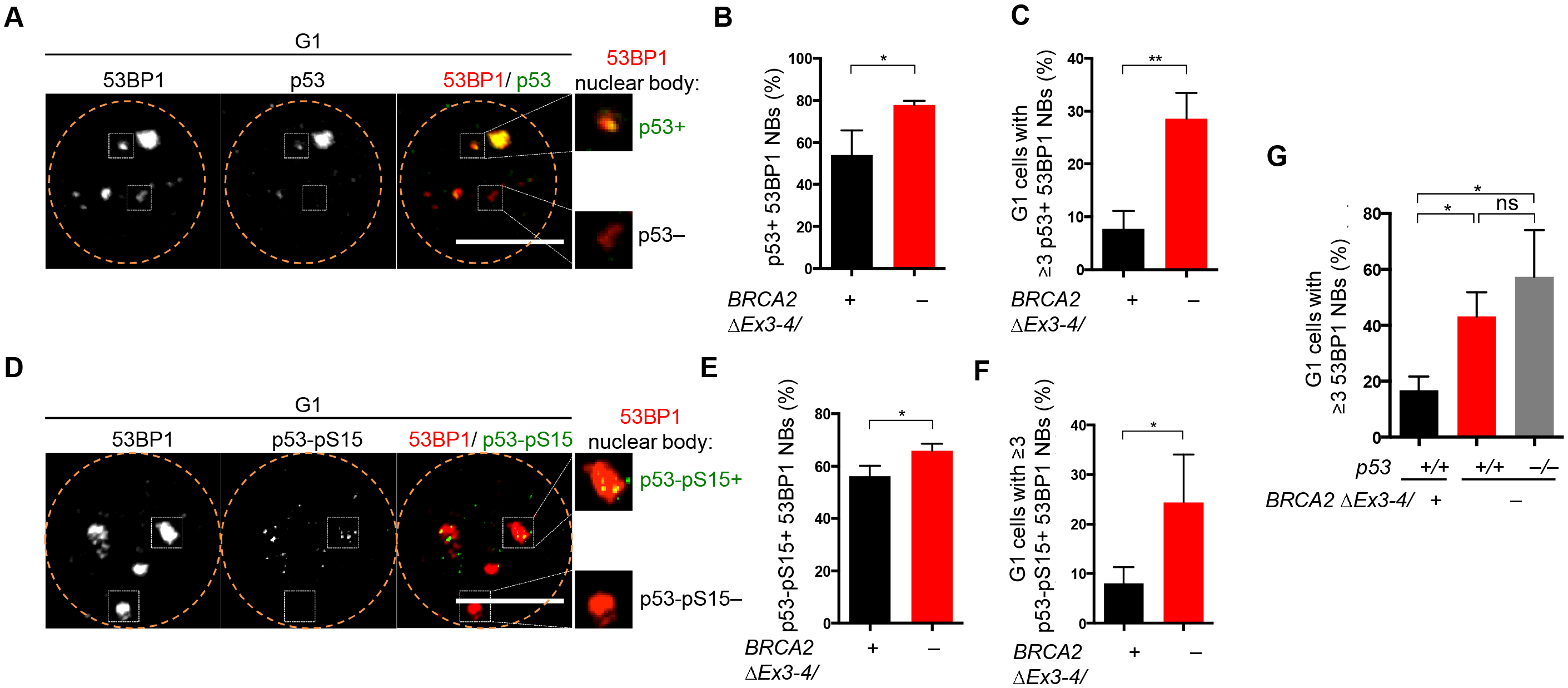
53BP1 nuclear body formation is associated with p53 activation. (A-F) Frequent 53BP1 nuclear body formation with p53 colocalization in G1 phase in BRCA2-deficient cells. Representative deconvolved images are shown with G1 nuclei (EdU-negative with 2N DNA content) outlined (A, D). Cells were pulse labeled with EdU for 30 min before IF. 53BP1 nuclear body formation and colocalization with p53 or p53-pS15 was quantified as in (B, C) and (E, F), respectively. Scale bars, 10 μm. (G) p53 loss does not significantly affect G1 53BP1 nuclear body formation in BRCA2-deficient cells. Error bars in this figure represent one standard deviation from the mean (s.d.). n=3. ns, not significant; ^*^, p<0.05; ^**^, p<0.01 (unpaired two-tailed t test).

53BP1 nuclear bodies could activate p53 at multiple levels [8], either directly through 53BP1-p53 interaction [40] or indirectly through the downstream DNA damage response. The underlying DNA lesions within the nuclear bodies may serve to activate signaling networks, including the p53 pathway. Indeed, 53BP1 nuclear bodies contain various components involved in the DNA damage response, including the master regulator ATM in its activated, phosphorylated form (pATM S1981) [30], which directly stabilizes and activates p53 [41]. Together, our results are consistent with a model that these nuclear compartments marked by 53BP1 serve as a signaling hub to recruit and activate p53. However, so far their relationship remains correlative and additional studies are needed to determine whether the relationship is causal and, if so, what are the mechanisms.

In addition to p53 activation, 53BP1 nuclear bodies may play other roles in BRCA2-deficient cells. For example, 53BP1 nuclear bodies sequester and shield the underlying sites of DNA damage [30]. Notably, removing 53BP1 dramatically enhances DNA break manifestation in the setting of CFS expression [30]. Thus, perturbing 53BP1 in BRCA2-deficient cells may similarly aggravate DNA break deprotection and thereby synergistically exacerbate genome instability phenotypes. This hypothesis, if found to be true, could potentially be exploited to develop novel synthetic lethal therapies for BRCA2-deficient cancers.

### Functional interplay between SMARCAL1 and BRCA2

Having uncovered some unusual aspects of BRCA2 deficiency, our BRCA2 conditional systems also afford the opportunity to dissect the contribution of two pathways, HR and FP. To this end, we developed multiple separation-of-function approaches to specifically perturb one pathway while keeping the other intact. These approaches converge on the conclusion that HR, rather than FP, plays the critical role in suppressing replication stress to support MCF10A cell survival [14]. Replication fork reversal has emerged in multiple studies as a new aspect dictating the outcome of FP versus fork degradation, suggesting that forks that have not reversed will not be degraded [8, 42]. We therefore examined the impact of fork reversal on replication stress by depleting SMARCAL1, a DNA translocase known to exhibit fork reversion activity [43, 44, 45]. In BRCA2-deficient cells, SMARCAL1 inactivation suppressed fork degradation (**Fig. 4A**), consistent with recent observations [11, 13] and the model that fork reversal acts as a prerequisite for nascent strand degradation (**Fig. 4B**). While FP was rescued, 53BP1 nuclear bodies were not significantly reduced in HU-treated BRCA2-deficient cells by SMARCAL1 depletion (**Fig. 4C**). Supporting these observations, inactivating ZRANB3, another DNA translocase that catalyzes fork reversal, also fails to abolish HU-induced chromosomal aberrations in BRCA2-deficient cells [12]. These results further corroborate our previous conclusions that FP is not critical for replication stress suppression.

**Figure 4.**
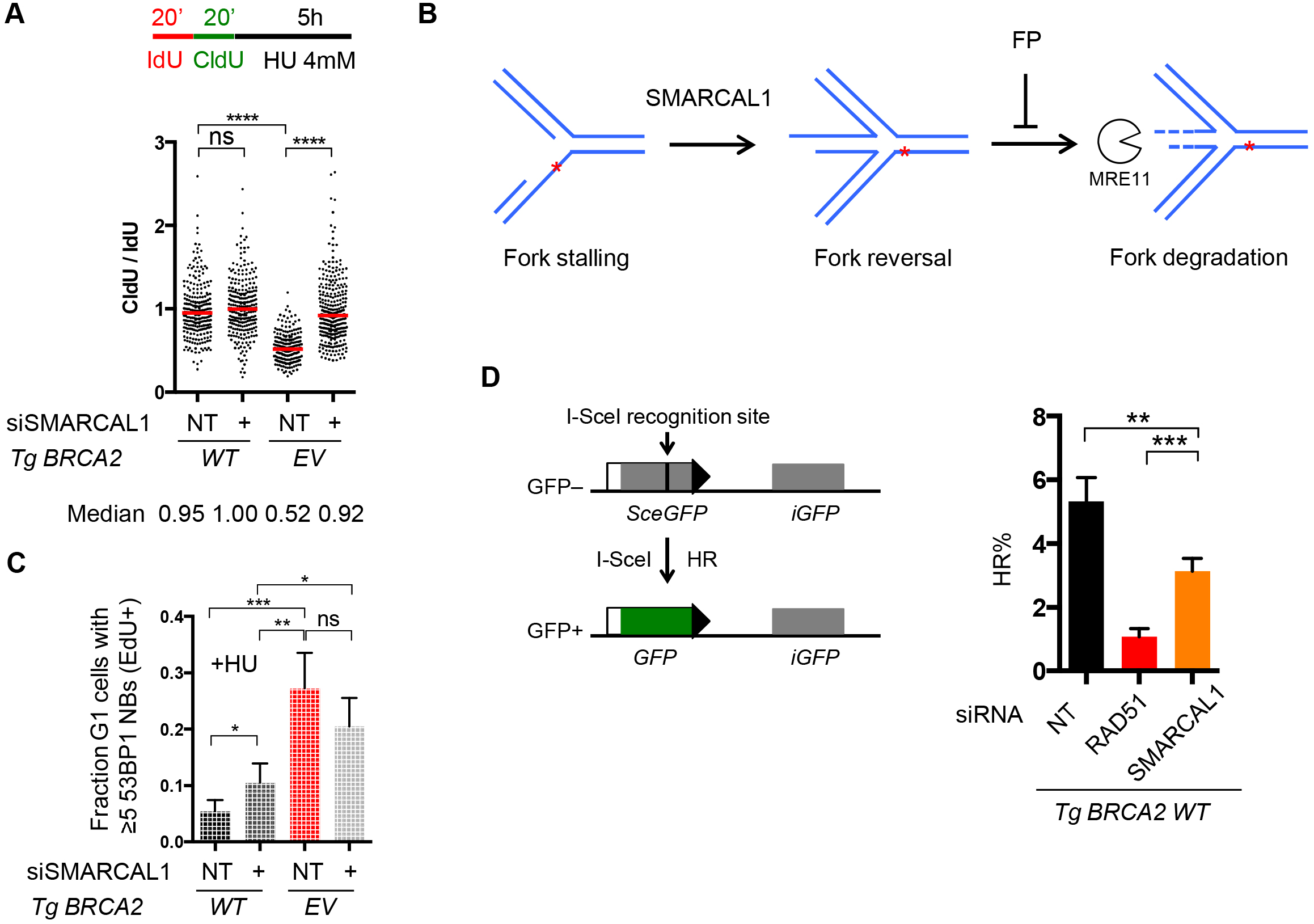
SMARCAL1 disruption in BRCA2-deficient cells restores FP but cells still maintain high 53BP1 nuclear body levels. (A) SMARCAL1 depletion restores FP to BRCA2-deficient cells. *BRCA2^∆Ex3-4/–^* cells expressing a WT BRCA2 transgene (*Tg BRCA2 WT*) or containing an empty vector (*EV*) were depleted of SMARCAL1 with siRNA. DNA fiber analysis was performed to quantify fork protection. Schematic of the experimental design is shown at top. Median CldU/IdU tract length ratios are indicated in the graph (red bars). Graphs represent the pooled results of >250 fibers per genotype from two independent experiments, analyzed by a two-tailed Mann-Whitney test. NT, non targeting siRNA. (B) A model depicting fork reversal as a step prior to fork degradation, based on recent studies. Upon fork stalling, replication forks are subject to reversal, a process mediated by DNA translocases including SMARCAL1. The resulting reversed forks serve as the entry point for subsequent nascent strand degradation, a reaction catalyzed by MRE11 nuclease and antagonized by the BRCA2-mediated FP pathway. (C) 53BP1 nuclear bodies are not significantly diminished in BRCA2-deficient cells by restoration of FP through SMARCAL1 depletion. *BRCA2^∆Ex3-4/–^* cells with or without the *BRCA2* transgene were depleted of SMARCAL1 and analyzed for HU-induced 53BP1 nuclear body formation, as in Fig. 2B. Statistical analysis was by an unpaired two-tailed t test. n=4. (D) SMARCAL1 depletion leads to an intermediate HR defect. A schematic of the DR-GFP reporter as a readout of HR activity is shown (left). *BRCA2^∆Ex3-4/–^;Tg BRCA2 WT* cells depleted of SMARCAL1 or RAD51 were infected with I-SceI-expressing lentivirus and the percent GFP+ cells was analyzed by an unpaired two-tailed t test (right). n≥3. Effective depletion of target proteins under the same conditions used in this study was previously confirmed by Western blot [14].

These data recapitulate previous results that SMARCAL1-mediated fork reversal plays a critical role in promoting nascent strand degradation upon replication stress [11, 13], adding to the expanding recent literature that implicates this process in the pathways involved in FP suppression [8]. FP restoration achieved by SMARCAL1 inactivation also represents an additional separation-of-function approach to study the contributions of the FP pathway in BRCA2- (and more generally HR-) deficient cells. The failure of FP to abolish the high replication stress found in BRCA2-deficient cells is in agreement with our previous findings obtained with orthogonal approaches [14]. Thus, although FP alone suffices to restore genome integrity in HR-deficient cancer cells under various genotoxic stresses [46], our results argue that FP plays a minor role in the relatively normal MCF10A cells. These apparent discrepancies could be due to distinct rate-limiting stresses that occur under different cellular contexts, e.g., under cancerous versus non-transformed states [8]. Future studies using additional non-transformed cell lines are needed to further validate this model.

Since fork reversal, either in excess or when limited, could cause DNA damage [44, 45, 47, 48], we examined whether SMARCAL1 impacts replication stress in BRCA2 WT cells. Interestingly, SMARCAL1 depletion by itself led to a modest enhancement of HU-triggered 53BP1 nuclear body formation (**Fig. 4C**). Given our previous findings that HR is critical to suppress replication stress [14], the observed induction in 53BP1 nuclear bodies is reminiscent of a role of SMARCAL1 in HR, as has been reported in the fly system [49]. To determine the HR function of SMARCAL1 in MCF10A cells, we employed the DR-GFP reporter that is stably integrated as a single copy to assay HR activity [50, 51] (**Fig. 4D**). Indeed, SMARCAL1 depletion led to a reduction in HR, although only to a moderate extent compared with that observed in cells depleted of the core HR component RAD51 [52] (**Fig. 4D**). Thus, SMARCAL1 depletion by itself exerts a small increase in 53BP1 nuclear body formation that may be associated with its minor role in HR. We conclude that SMARCAL1 plays a minor role in suppressing replication stress when BRCA2 is present. In sum, our previous [14] and current results are all consistent with a model that HR plays a critical role in suppressing replication stress leading to 53BP1 nuclear bodies, whereas neither protection nor regression of replication forks substantially contributes to this process.

### Concluding remarks

We have previously shown that BRCA2 suppresses DNA replication stress through HR [14]. We have now expanded on these findings, investigating three aspects of the replication stress triggered as a consequence of BRCA2 inactivation. First, we show that BRCA2-deficient cells are sensitized to two types of replication stress, those induced by HU and by low-dose APH. Second, 53BP1 nuclear bodies spatially link replication stress to downstream p53 activation in G1 phase. Finally, disrupting fork reversal by SMARCAL1 depletion ameliorates the FP defect in BRCA2-deficient cells, but does not restrain 53BP1 nuclear body formation, further corroborating our previous model that FP plays a minor role in suppressing replication stress in MCF10A cells.

## Materials and methods

### MCF10A Cell culture and chemicals

*BRCA2^fl/+^*, *BRCA2^fl/-^*, *BRCA2^fl/-^;p53^-/-^* and *BRCA2^fl/-^;Tg BRCA2 WT/EV* cells were previously generated [14] from MCF10A DR-GFP cells [50]. *BRCA2^∆Ex3-4/+^* and *BRCA2^∆Ex3-4/–^* cells are generated from the pool of *BRCA2^fl/+^* and *BRCA2^fl/–^* cells respectively, 3-6 days after lentiviral transduction of self-deleting Cre recombinase, as previously described [14].

MCF10A cells were cultured in DME-HG/F-12 supplemented with 5% horse serum, 20 ng ml^−1^ epidermal growth factor, 0.5 mg ml^−1^ hydrocortisone, 10 μg ml^−1^ insulin, 100 ng ml^−1^ cholera toxin, and 1% penicillin–streptomycin.

Aphidicolin (0.1 μg ml^-1^; A4487, Sigma), hydroxyurea (4 mM; H8627, Sigma), IdU (50 μM; I7125, Sigma), CldU (50 μM; C6891, Sigma) were used for treatment as indicated.

### Plasmid construction and gene targeting with CRISPR-Cas9

Hyg-targeting was performed as previously described [14]. Briefly, donor plasmid contains homology arms that are 700-900 bp in length. Wild-type Cas9-expression plasmid (Addgene plasmid # 41815) [53] (**Fig. 1**) or paired nickases (Cas9 H840A) [54, 55] (**Fig. S1**) and BRCA2 sgRNA(s), which were cloned into sgRNA backbone (Addgene plasmid # 41824) [53], were used together with the donor plasmid. Transfection by electroporation (Gene Pulser II, Bio-Rad; 350 V, 1000 μF) was performed before selection for hygromycin (50 μg ml^−1^) resistance.

For genotyping PCR, trypsinized cell suspensions were mixed with PCR grade water and heated at 100 °C for 5 min before PCR analysis. Oligos 1 (forward) and 2 (reverse) were used to detect the Hyg-targeted allele. Oligos 1 (forward) and 3 (reverse) were used to detect the allele that had not been targeted by the Hyg cassette (i.e., the untargeted allele). Oligo 4 was used to characterize the untargeted allele by Sanger sequencing.

sgRNA target sequences

For *BRCA2* exon 2 G allele targeting:

G allele-specific sgRNA: AGACTTATTTACCAAGCAT

For *BRCA2* exon 2 A allele targeting:

A allele-specific sgRNA: AGACTTATTTACCAAACAT

Another *BRCA2* exon 2 sgRNA (not allele specific, used together with the A allele-specific sgRNA for paired nickase strategy [54]): GCCTCTCTTTGGATCCAAT

DNA oligo sequences:

Oligo 1: GCTTCTGAAACTAGGCGGCAG

Oligo 2: ATATCCACGCCCTCCTACATCG

Oligo 3: AATGTTGGCCTCTCTTTGGATC

Oligo 4: TCACTGGTTAGCGTGATTGAAAC

### Lentiviral transduction

Lentivirus was produced following standard methods. Briefly, HEK293T cells were co-transfected with a lentiviral vector, VSV-G expression plasmid and psPAX2 by Lipofectamine 2000 (11668027, Thermo Fisher Scientific) following the manufacturer’s instructions. Virus-containing supernatants were collected and 0.45-μm filtered 48 and 72 h after transfection. Infections of MCF10A cells were performed in the presence of 8 μg ml−1 polybrene (TR-1003- G, EMD Millipore).

### RNA interference

Transient depletion of SMARCAL1 and RAD51 was achieved by transfecting small interfering RNA (siRNAs) using Lipofectamine RNAiMax (13778075, Thermo Fisher Scientific), according to the manufacturer’s instructions, 24 h or 48 h after Cre infection. siRNAs were purchased from vendors: RAD51 (Qiagen, SI03061338), SMARCAL1 (SMARTpool, Dharmacon, M-013058-01-0005).

### Immunofluorescence and microscopy

Immunofluorescence was performed as described [14]. Briefly, cells were seeded on Nunc^™^ Lab-Tek^™^ II CC2^™^ Chamber Slides (12-565-1, Thermo Fisher Scientific), treated as indicated before fixation with 2% paraformaldehyde and permeabilization in blocking buffer (0.1% Triton-X and 1% BSA in PBS). Where indicated, EdU was detected using Click-iT^®^ Plus EdU Alexa Fluor^®^ 647 Imaging Kit (C10640, Thermo Fisher Scientific). Primary antibodies and secondary antibodies were sequentially applied. Slides were mounted with Prolong with DAPI (P36935, Thermo Fisher Scientific).

For APH/HU-induced 53BP1 nuclear body analyses (**Fig. 2**), images were acquired under Axio2 microscope (Zeiss). For 53BP1-p53/p53-pS15 colocalization analyses (**Fig. 3**), images were acquired under DeltaVision Elite microscope (GE Healthcare). Deconvolution was carried out with z stacks acquired with 0.2 μm spacing using enhanced ratio method, and projected based on maximum intensity on a DeltaVision Image Restoration System (GE Healthcare). Image analysis was performed using high-content image-based cytometry methods as previously described [14, 56], using FIJI (ImageJ).

Primary antibodies were used as follows: 53BP1 (1:1000; 612522, BD Biosciences), 53BP1 (1:1000; NB100-304, Novus), cyclin A (1:1000; sc-751, Santa Cruz Biotechnology), p53 (1:1000; Santa Cruz, sc-98), p53-pS15 (1:1000; CST, 9284). Secondary antibodies were used as follows: anti-mouse Alexa Fluor^®^ 488, anti-rabbit Alexa Fluor^®^ 488, anti-rabbit Alexa Fluor^®^ 568, anti-mouse Alexa Fluor^®^ 594, anti-rabbit Alexa Fluor^®^ 594 (1:1000; Thermo Fisher Scientific).

### DNA fiber assay

DNA fiber assay for FP analysis was performed as previously described [14]. Briefly, cells were pulse labeled with 50 μM IdU and 50 μM CldU, treated with 4 mM HU and lysed in lysis buffer (0.5% SDS, 200 mM Tris-HCl pH 7.4, 50 mM EDTA) before spreading on microscope slides followed by fixation (methanol/acetic acid at 3:1 ratio by volume). DNA was denatured in 2.5 M HCl, blocked in blocking buffer (10% goat serum and 0.1% Triton-X in PBS), followed by immunostaining with primary antibodies (anti-CldU, 1:75, MA1-82088, Thermo Fisher Scientific; anti-IdU, 1:75, 347580, BD Biosciences) and secondary antibodies (anti-rat Alexa Fluor^®^ 488 and anti-mouse Alexa Fluor® 594, 1:250, Thermo Fisher Scientific). Slides were mounted in Prolong with DAPI (P36935, Thermo Fisher Scientific) before image acquisition under DeltaVision Elite microscope (GE Healthcare). Images were analyzed with FIJI (ImageJ) software.

### HR assay

Cells were infected with an I-SceI-expressing lentivirus [14]. 48 h later, HR was quantified by measuring the fraction of GFP-positive cells using flow cytometry (Becton Dickinson FACScan) and FlowJo software.

## Acknowledgements

We thank members of the Jasin lab for discussions and suggestions. We thank Scott Keeney and Prasad Jallepalli for discussions. This work was supported by an Olayan Fellowship (W.F.), MSK Cancer Center Support Grant/Core Grant P30 CA008748, and R01 CA185660 (M.J.).

---

**Figure S1. Gene targeting at the *WT* allele of *BRCA2^fl/+^* cells.**

Schematic for generation of *BRCA2^fl/–^* cells described in [14]. The Hyg cassette was designed to be specifically targeted to the *BRCA2 WT*, but not the *fl* allele. Unlike in **Fig. 1**, the target allele (*WT*) contains the A nucleotide SNP, so an A allele-specific sgRNA was designed to achieve the Hyg cassette targeting (with a paired nickase strategy, see methods). Filled triangles, loxP sites; open circle, FRT site. Other symbols are as described in the legends of **Fig. 1A**.

